# The topography of DNA replication origins in Eukarya: GGN clusters, landmark nucleosomes, CDC6 and G4 structures

**DOI:** 10.1101/2024.01.04.574144

**Authors:** Ugo Bastolla, Irene Gómez-Pinto, Zaida Vergara, María Gómez, Carlos González, Crisanto Gutiérrez

**Affiliations:** Centro de Biología Molecular “Severo Ochoa” CSIC-UAM Cantoblanco, 28049 Madrid, Spain; Instituto de Quimica Fisica Blas Cabrera (CSIC), Madrid, Spain

## Abstract

**Background:** We recently identified tandem repeats of GGN triplets as a sequence motif strongly enriched in the DNA replication origins (ORIs) of the *Arabidopsis thaliana* genome, where it correlates with the ORI strength quantified through short nascent strands sequencing.

**Results:** Here we show that clusters of four or more GGN repeats separated by short tracts (<7 nucleotides) are present in more than 65% ORIs of six model eukaryotic organisms: *Leishmania major*, *Arabidopsis thaliana*, *Caenorhabditis elegans*, *Drosophila melanogaster*, *Mus musculus* and *Homo sapiens*. The measured percentages vary with the experimental technique adopted for ORI determination. For all studied techniques and organisms, ORIs are significantly prone to be located within or near strong GGN clusters, although these are neither necessary nor sufficient for ORI activation. Interestingly, for same experimental technique, the association between ORIs and GGN clusters was stronger al later developmental stages for all organisms for which we could perform this comparison.

In accordance with a biophysical model of nucleosome positioning, the GGN clusters strongly favor nucleosome occupancy. At the same time, both GGN clusters and ORIs occur frequently within 1kb from nucleosome-depleted regions (NDR). We propose a structural model based on chromatin secondary structure in which the NDR and the well-positioned nucleosome at the GGN are close in space, which may favor functional interactions. We hypothesize that the presence of GGN at ORIs arose in part for promoting the above nucleosome organization Moreover, at least in *Arabidopsis*, the replication protein CDC6 is very strongly associated with GGN clusters.

NMR experiments showed that clusters of at least four GGN can form G-quadruplex (G4) *in vitro.* Our data support the view that GGN clusters are formed through the interplay of mutational processes (GC skew at ORIs plus triplet expansion), and that similar mutational processes, i.e. AT skew at transcription start sites (TSS) and ORIs, might facilitate the formation of the NDR, thus favoring the evolvability of chromatin.

**Conclusions:** GGN clusters are easily evolvable sequence motifs enriched at ORIs of eukaryotic genomes. They favor G4 secondary structure and a nucleosome organization that may unify the apparent discrepancy between ORIs of higher eukaryotes and yeast.

## Background

Genome replication is crucial for transferring the genetic information to the daughter cells during the cell cycle. In eukaryotic organisms, one of its complexities derives from the need to start DNA synthesis at thousands of genomic locations (DNA replication origins, ORIs) in a coordinate manner during S-phase. Despite extensive investigation, the sequence and structural determinants of ORIs in most Eukarya are not yet fully understood. Genome-wide ORI mapping in several of these organisms has revealed that ORI midpoints contain G-rich sequences. These include mammal, fly, worm and plant cells (Costas et al. 2011; Picard et al. 2014; Cayrou et al. 2015; Comoglio et al. 2015; Fu et al. 2015; Lombraña et al. 2016; Pourkarimi et al. 2016; Rodriguez-Martinez et al. 2017; Vergara et al. 2017; Sequeira-Mendes et al. 2019; Akerman et al. 2020). Several articles reported an association between ORIs and atypical DNA secondary structures formed by stacked arrangements of planar G-tetrads stabilized by Hoogsteen hydrogen bonding, called G-quadruplexes (G4) (Hansen et al., 2010; Cayrou et al., 2011; Besnard et al., 2012; Gindin et al., 2014; Valton et al., 2014; Smith et al., 2016; Prorok et al., 2019). While the role of G4s in ORI activity is still a matter of debate (Valton and Prioleau 2016; Akerman et al., 2020), a recent pre-print demonstrated that engineering dimeric G4 motifs in the same DNA strand is sufficient to induce the formation of a minimal ORI (Poulet-Benedetti et al., 2023).

We recently showed that ORIs in the model plant *Arabidopsis thaliana* are associated with several repeats of GGN triplet motifs, where N is any nucleotide (Sequeira-Mendes et al. 2019). Here we set out to investigate in a comparative and comprehensive manner the propensity of clusters of GGN triplets to associate with eukaryotic ORIs and the possible evolutionary mechanisms through which they could arise. To this end, we examined ORIs of six eukaryotic genomes: *Homo sapiens* (Picard et al., 2014; Fu et al. 2015; Petryk et al. 2016; Langley et al. 2016), the mouse *Mus musculus* (Almeida et al., 2018), the fly *Drosopila melanogaster* (Comoglio et al., 2015), the worm *Coenorabditis elegans* (Pourkarimi et al., 2016), the parasite *Leishmania major* (Lombraña et al., 2016) and the plant *Arabidopsis thaliana* (Costas et al. 2011; Sequeira-Mendes et al. 2019; Wheeler et al. 2020). We found that, in almost all cases, ORIs are enriched in GGN clusters and, conversely, GGN clusters show a marked propensity to appear at ORIs, although the presence of a GGN cluster is neither necessary nor sufficient for ORI specification. The only exceptions to this association are *C. elegans* pre-gastrula embryos, but not post-gastrula ones, and initiation regions of *A. thaliana* (Wheeler et al. 2020) but not ORIs of the same organism determined through other methods. In general, we found a strong influence of the experimental method of ORI determination, following known experimental biases that we discuss below. However, we also found significant association for methods that present opposite bias, such as short nascent strand sequencing (SNS-seq) and labeling with thymidine analogs (BrdU-seq), which indicates that these biases cannot be the only cause of the reported association. Strikingly, the association between GGN and ORIs is stronger at later developmental stages for all organisms for which we could perform this comparison, independent of the experimental approach. At least in *A. thaliana*, GGN clusters are also associated with nucleosomes and, very strongly, with the replication protein CDC6, which may explain their association with ORIs. We complemented these computational analyses with circular dichroism and NMR experiments, demonstrating that four consecutive triplets of GGA or GGT repeats are able to form G4 secondary structures *in vitro*. Interestingly, the G4-stability ranking of the third base of the GGN recapitulates the corresponding observed frequency in the studied genomes, consistent with selective pressure favoring G4 stability at ORIs. Finally, our data support the hypothesis that repeated triplets of nucleotides, including GGN and AAN, arise in evolution through a process of triplet expansion, superimposed with and confounded by smaller insertions and/or deletions, which helps explaining ORI evolvability.

## Results

### DNA replication origins (ORIs) are enriched in GGN clusters

We examined ORIs of six eukaryotic genomes mapped through different techniques (see Supplementary Table 1). (1) Sequencing of short nascent strands (SNS-seq) purified through sucrose size fractionation and extensive digestion with lambda exonuclease (λ-exo): *H. sapiens* (Picard et al., 2014, Fu et al. 2015), *M. musculus* (Almeida et al., 2018), *D. melanogaster* (Comoglio et al., 2015), *L. major* (Lombraña et al., 2019) and *A. thaliana* (Sequeira-Mendes et al. 2019). (2) Sequencing of Okazaki fragments (OK-seq): *H. sapiens* (Petryk et al. 2016) and *C. elegans* (Pourkarimi et al., 2016). (3) Labeling with the thymidine analog BrdU followed by density fractionation and sequencing (Brdu-seq): *A. thaliana* (Costas et al. 2011). (4) Labeling with the thymidine analog EdU followed by two-parameters cell sorting and immunoprecipitation (EdU-seq): *A. thaliana* (Wheeler et al. 2020). (5) Cell-free labelling of early-replicating DNA followed by immunoprecipitation (Ini-seq): *H. sapiens* (Langley et al. 2016). Considering these different methodologies allowed us to investigate the possible biases derived from each experimental procedure.

The number of identified ORIs vary widely depending on the organism, cell type or developmental stage and experimental technique, ranging from 12 ORIs *per* mega-base (*H. sapiens* SNS-seq from Picard et al., 2014) to 209 ORIs *per* mega-base (*L. major* SNS-seq) (Figure 1A). The mean ORI length reported also varied broadly, from the high resolution of the SNS-ORIs of *A. thaliana* (0.48 kb) and *M. musculus* (0.63 or 0.67 kb depending on cell type), to the broad initiation regions (IR) of *H. sapiens* detected through OK-seq (34 kb) (Figure 1B). Finally, the total length of the genome covered by the mapped ORIs ranged from 1.1% (*A. thaliana* SNS-seq) to 20% of the genome (*L. major* SNS-seq) (Figure 1C). There is only moderate coincidence between ORIs determined in different cell types of the same organism, and ORIs determined with different techniques often coincide less than or same as expected by chance. However, the *H. sapiens* ORIs determined through Ini-seq (Langley et al., 2016) and those determined through SNS-seq by Fu et al. (2015), and the *A. thaliana* ORIs determined through SNS-seq and BrdU-seq (Sequeira-Mendes et al., 2019; Costas et al., 2011) coincide with each other much more than expected by chance (Supplementary Fig. S1).

**Figure 1:**
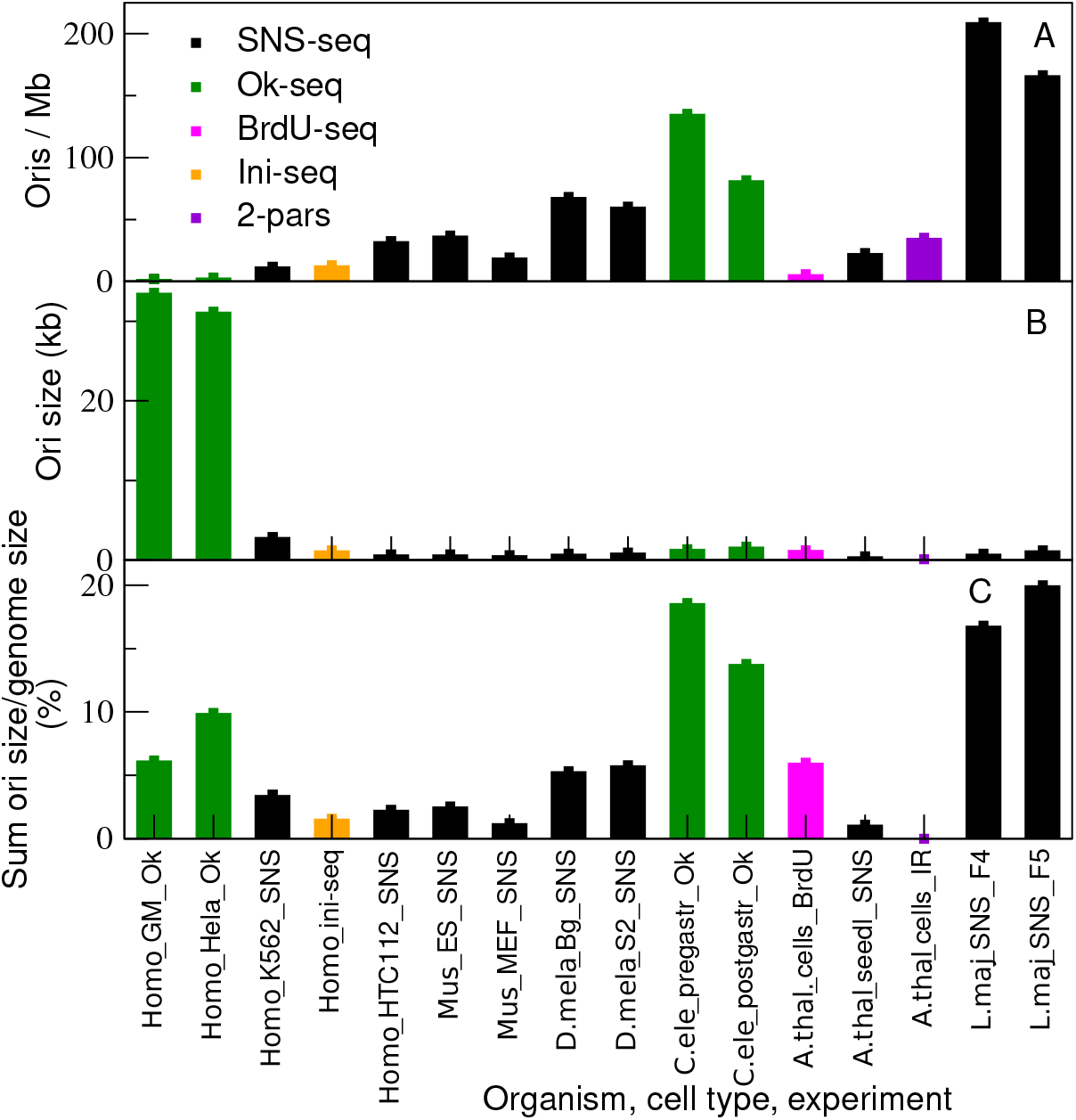
Properties of the ORI sets studied in this work. A: Number of ORIs per mega-base (Mb); B: Average length of the ORIs reported in the bed file (kb). C: Total length of the reported ORIs with respect to genome size (%). Note that the same organism may appear in sets from different cell types, developmental stages or experimental techniques.

Using our program *GGN_cluster*, we retrieved from all genomes clusters of at least *n* consecutive GGN motifs (GGN >= *n*) separated by at most *l<=L* other nucleotides (see Methods for details). As expected, the number of clusters increases with the G+C content of the genome, which is 35% (*C. elegans*), 36% (*A. thaliana*), 38% (*M. musculus*), 40% (*H.sapiens*), 42% (*D. melanogaster*) and 58% (*L. major*), strongly decreases with the minimum number of GGN triplets in the cluster, *n,* and strongly increases with the maximum separation *L* (Supplementary Fig. S2). The bed files with the coordinates of the GGN clusters for different parameters *n* and *L* are included in the additional file GGN_clusters.zip.

We then analyzed the propensity of ORIs to associate with GGN clusters. Figure 2 represents the mean Z score of ORIs versus the distance from the center of the clusters with GGN>=8 and L=6, which is a typical case. Error bars represent the standard errors of the mean, and allow to visually assess the significance of the association. We found that the strength of the association depends on the organism, the cell type and, to large extent, the experimental method for ORI determination, which present known biases. We address the latter by discussing the cases of *Arabidopsis thaliana* and *Homo sapiens* ORIs, for which we can compare different methods.

**Figure 2:**
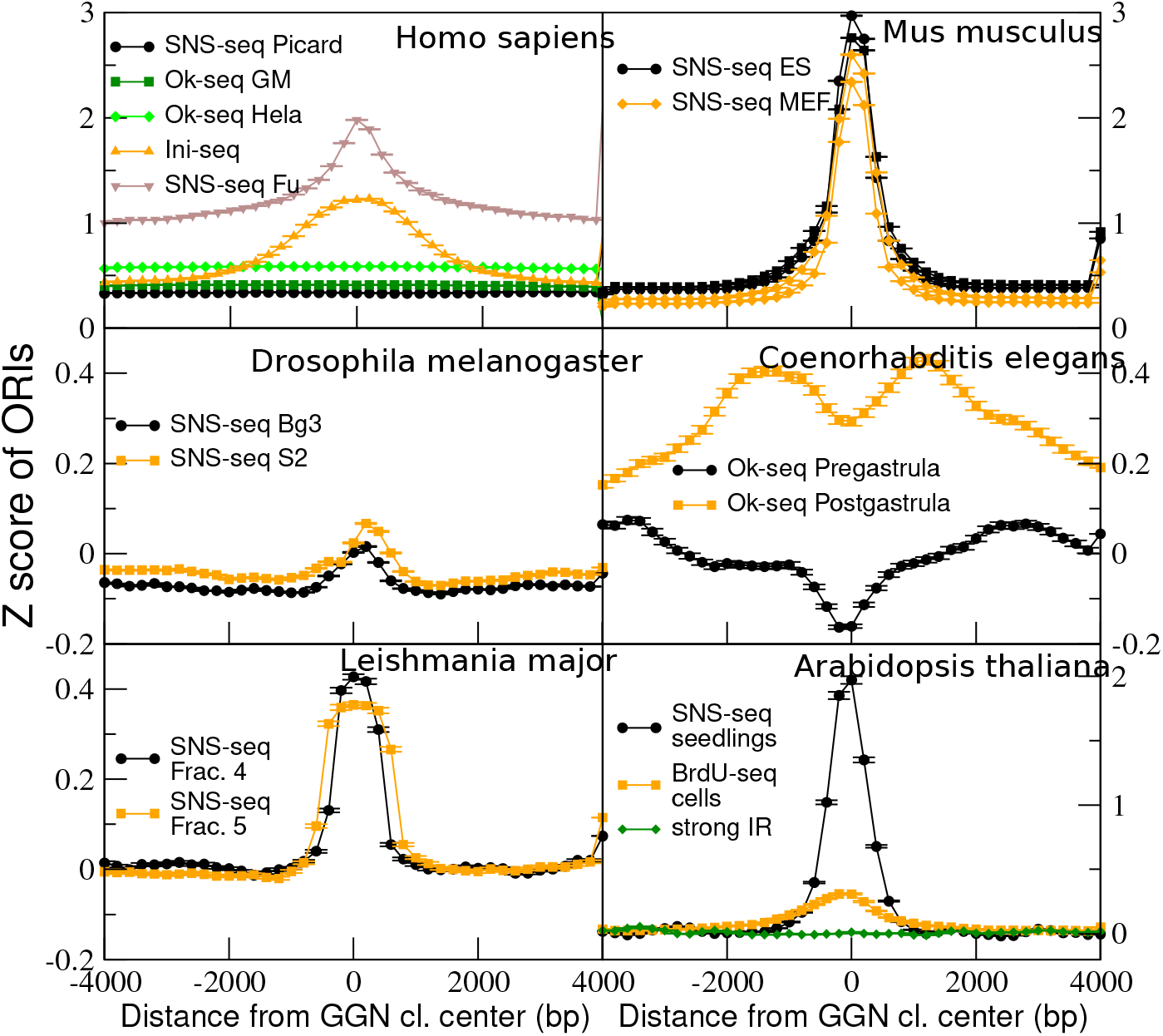
Z score of ORI score as a function of the distance from the center of GGN clusters with GGN≥8 and L≤6.

Digestion with λ-exo, which is a crucial step for identifying ORIs by SNS-seq, is biased towards GC-rich sequences (Foulk et al., 2015) and it may account for the GC propensity observed in several ORI sets. This bias can be reduced by using as control genomic sequences also subject to λ-exo digestion (Sequeira-Mendes et al. 2019). On the contrary, ORIs determined through labeling with thymidine analogs such as BrdU and EdU are biased towards AT-rich sequences. This bias can be reduced weighting the control with the number of A or T bases in each sequence. To this end, we reanalyzed ORIs mapped by BrdU-seq of Costas et al. (2011) using our algorithm Zpeaks (Sequeira-Mendes et al., 2019) with AT-weighted control, and we identified 5800 ORIs with mean size of 1.2 kb versus 1500 ORIs with size 1.4 kb in the original data set, see Figure 1. Their properties are very similar to those of the 2700 SNS-ORIs of *Arabidopsis* identified by Sequeira-Mendes et al. (2019), with which they share 756 ORIs, 3 times more than expected by chance (P<0.005), an overlap in a similar range when comparing other ORI sets in the same organism determined through the same method (Supplementary Fig.S1). Despite the inherent AT bias of the mapping method, the BrdU-ORIs are strongly GC rich, as it is also the original dataset by Costas et al. (2011).

The *A. thaliana* ORIs determined by SNS-seq preceded by λ-exo digestion are strongly associated with GGN clusters, but also BrdU-ORIs show a significant association. The difference between the two sets (Z score of 2 for SNS-seq versus 0.35 for BrdU-seq, Figure 2, lower right panel) illustrates their differential biases towards G+C rich and A+T rich sequences, respectively. The association is in both cases highly significant due to the large number of examined ORIs.

In contrast, the IRs of *A. thaliana* determined by Wheeler et al. (2020) do not show any enrichment at GGN clusters (Fig.2), even when we apply the AT-weighted control. We found that these IRs have a strong propensity to occur at Polycomb-repressed chromatin states and avoid active chromatin states (Supplementary Fig. S3D). This result is puzzling, because ORIs determined by the other two approaches tend to occur in active chromatin states (Supplementary Fig. S3B-C). In general, these IRs displayed contrasting properties to the other two sets of *A. thaliana* ORIs; they do not colocalize with transcribed regions, they tend to be located at nucleosome depleted regions, they are GC-poor, and, most importantly, they are not enriched in the pre-replication complex protein CDC6 (Supplementary Fig. S4). These properties are different from those reported in most other eukaryotic organisms, in which ORIs are strongly correlated with transcriptionally active chromatin (Picard et al. 2014; Cayrou et al. 2015; Comoglio et al. 2015; Lombraña et al. 2016; Pourkarimi et al. 2016; Rodriguez-Martinez et al. 2017; Akerman et al. 2020; Jodkowska et al., 2022). We hypothesized that these peculiar properties may derive from the non-uniformity of EdU incorporation throughout the genome or their large size. We reanalyzed the data by Wheeler et al. (2020) to take into account the possibility that EdU incorporates non-uniformly in the genome, and we identified a new set of putative ORIs with properties very different from those of the original set. For reasons of space, we will present this analysis elsewhere and we will not examine further in this study the ORIs reported by Wheeler et al. (2020).

For *H. sapiens*, we considered five sets of ORIs determined with three different methods: SNS-seq (Picard et al. 2014, Fu et al. 2015), Ini-seq (Langley et al. 2016) - mapping method that do not use λ-exo digestion -, and IR determined through EdU labelling followed by Ok-seq (Petryk et al. 2016). In spite of the different biases of each mapping technique, all sets associate with GGN clusters with a Z score significantly larger than zero (Fig.2 upper left). Ini-seq ORIs present a stronger enrichment at the GGN center than the SNS-ORIs from Picard et al. (2014), ruling out the putative bias for GC-rich sequences as the only explanation for this association. To note, the IR from Petryk et al. (2016) and the SNS-ORIs from Picard et al. (2014), which span several kb, are uniformly distributed along more than 4 kb around the GGN center, while the SNS-ORIs from Fu et al. (2015) and the Ini-seq ORIs from Langley et al. (2016), peak at the GGN center. In the following sections, we only consider these later two sets of human ORIs because of their better resolution (Supplementary Figure S1).

The strongest of all examined associations (Z score=3; Fig. 2, upper right plot) occurs at the SNS-seq *M. musculus* ORIs (Almeida et al., 2018); while *L. major* ORIs (Lombraña et al., 2016), also determined by SNS-seq, display an intermediate Z score of approximately 0.4 (Fig. 2, lower left). For the same organism and mapping approach, association scores and patterns differ between cell types. The association is stronger for *C. elegans* post-gastrula embryos than pre-gastrula embryos (Z score of 0.4 and −0.2, the latter lower than expected by chance, Fig.2 middle-right plot), both determined through OK-seq (Pourkarimi et al., 2016). Similarly, it is stronger for *D. melanogaster* S2 cells than for Bg3 cells (Z score of 0.1 and 0.03, being the latter as expected by chance, Fig.2 middle left plot), both determined through SNS-seq (Comoglio et al., 2015). The results were very similar when considering GGN clusters with GGN >= 4 and L=0 (Supplementary Fig. S5). Altogether, these analyses indicate that ORIs are prone to be located at or very near to GGN clusters in the genome of several organisms, despite the marked differences in the number of identified ORIs, methodology of ORI mapping, and GC richness.

We complemented this analysis by evaluating the propensity of GGN clusters to occur close to ORIs, illustrated by the mean Z score of GGN clusters versus the distance from the ORI midpoints (Supplementary Figures S6, GGN >= 8 L=6 and S7, GGN >=4, L=0). GGN clusters are significantly prone to be located near ORIs in all organisms, but the Z scores are lower than for the propensity of ORIs to be located at GGN shown in Figure 2. Then we examined the fraction of ORIs containing GGN clusters as a function of the minimum number of GGN triplets in the cluster (*n*) and the maximum number of bases between consecutive triplets (*L*) for different *L* values (Figure 3). A large fraction of ORIs contain clusters with at least 4 GGN even with the strict threshold *L*=0. Most overlaps are significantly larger than expected by chance at the 1% level (noted by ++ in Figure 3; see Methods for details). The smallest fractions were found for *C. elegans* (7.5% for pre-gastrula and 9.4% for post-gastrula ORIs, respectively), less than expected by chance. For clusters with GGN≥4 and the less stringent threshold *L ≤* 3, the fraction of ORIs that overlap with GGN clusters increases considerably, ranging from 100% in *L. major* to ∼30% in *C. elegans* (Figure 4). Finally, allowing *L ≤* 7 bases between consecutive GGN motifs, the fraction of ORIs become 74-81%, 73-82% and 65% for *D. melanogaster*, *A. thaliana* and *C. elegans* genomes, respectively, the latter again smaller than expected. We conclude that most ORIs, although not all of them, contain one or more clusters with at least four GGN motifs. One interesting observation from this analysis is that, despite the strong dependence on the G+C content, the fraction of ORIs that contain GGN clusters can be quite high even in GC-poor genomes such as *A. thaliana* and *M. musculus.* The enrichment (ratio between observed and expected number of ORIs containing different amounts of GGN triplets) is significantly ≥1 for most systems and GGN clusters (Supplementary Fig. S8).

**Figure 3:**
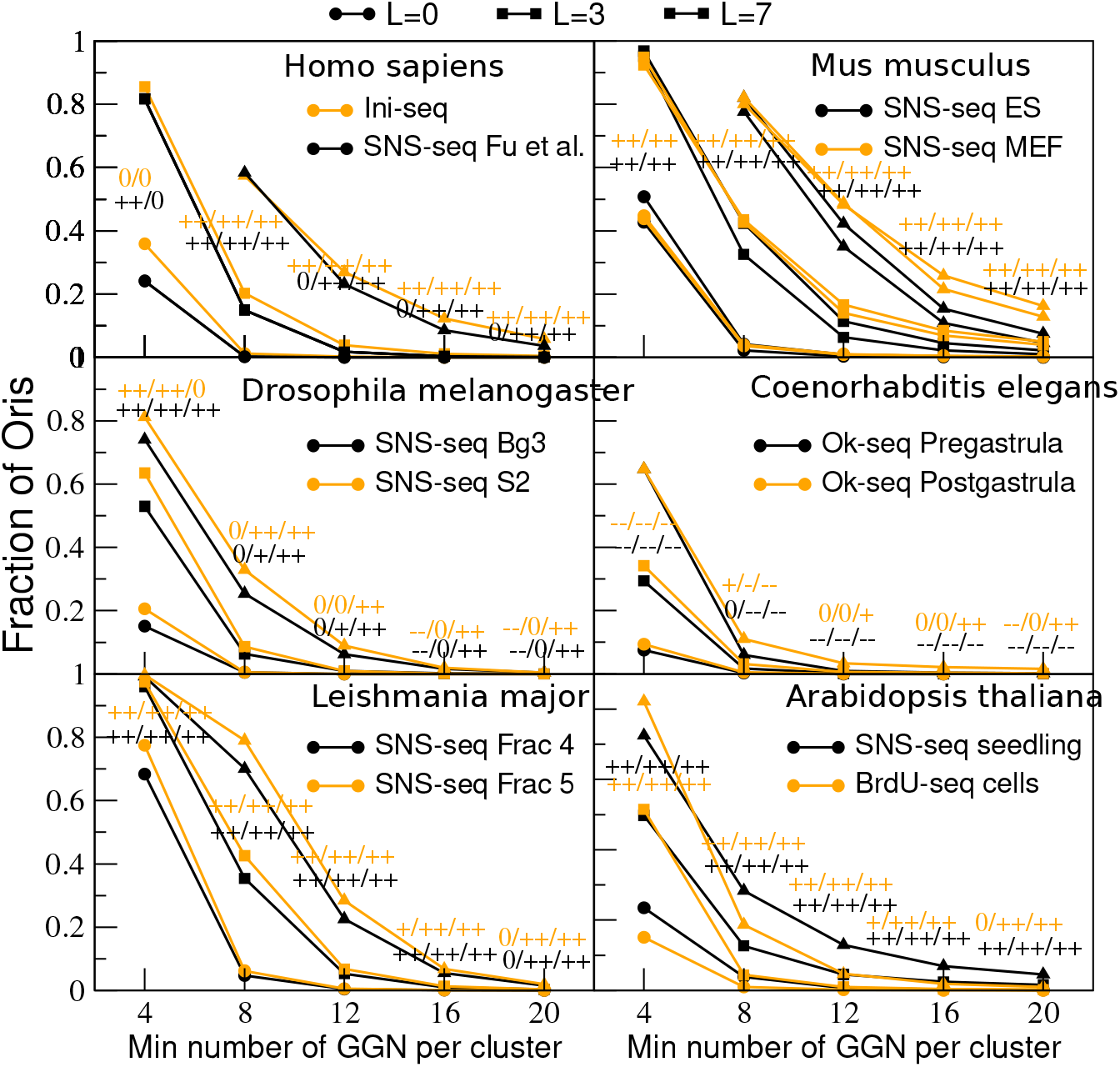
Fraction of ORIs containing a cluster with at least *n* GGN motifs (horizontal axis). The symbols represent the maximum allowed separation between consecutive GGN motifs in the cluster: *L*=0 (circles), *L=*3 (square) and *L=*7 (triangles). Colors represent the cell types (black for earlier in development and orange for later stage), the experimental fraction for *L. major* or the experimental technique for *A. thaliana* (black: SNS-seq; orange: BrdU-seq). Significance is indicated with the symbol ++ (--) represent overlaps that are significantly larger (smaller) than expected by chance at the 0.01 level, + (-) represent significance between the 0.05 and the 0.01 level, and 0 represents no significance. Each line, such as ++/++/++, represents the significance of the *L*=0, *L=*3 and *L=*7 clusters, respectively.

**Figure 4:**
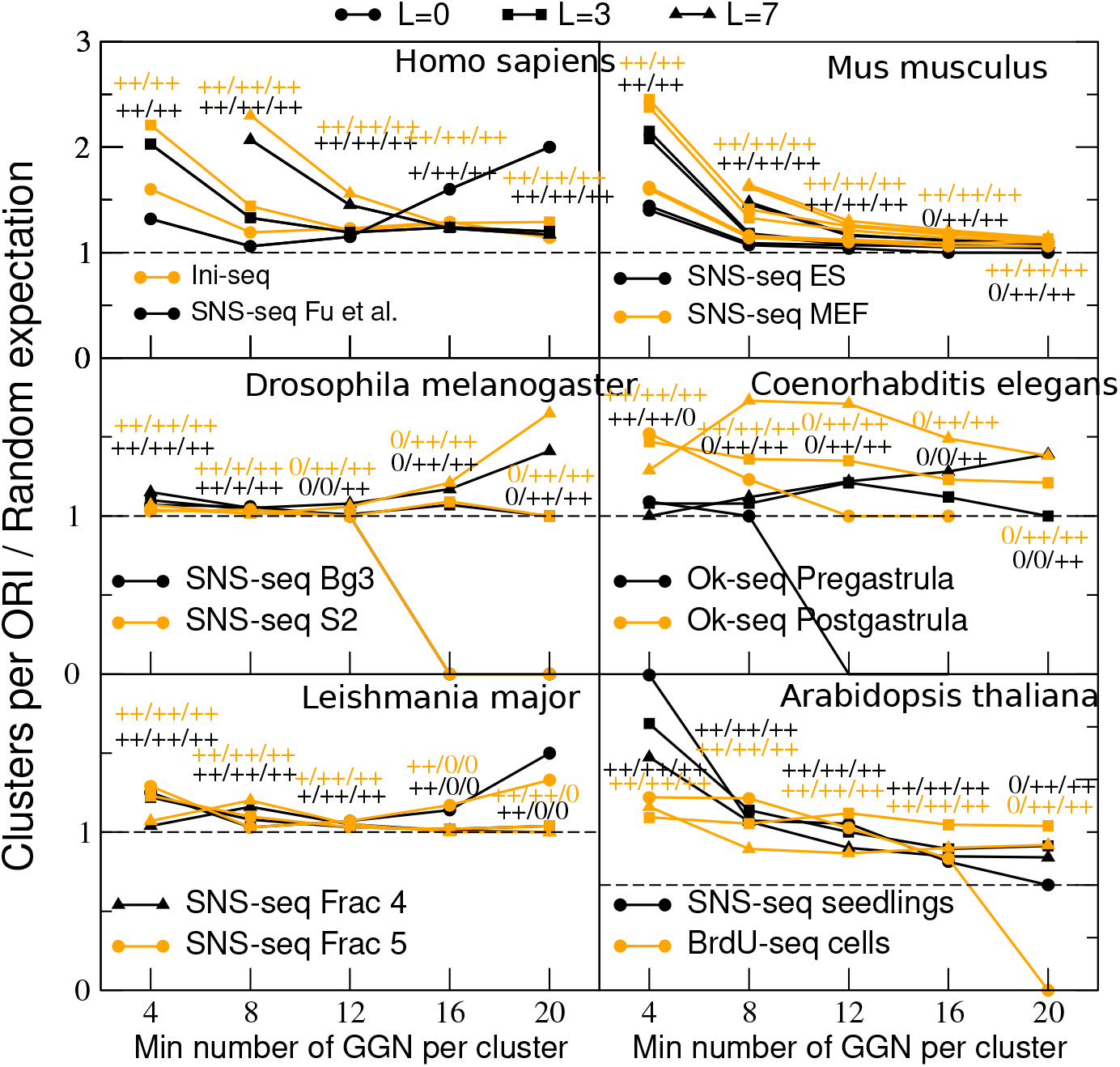
Enrichment of number of clusters per ORI. For ORIs that contain at least one GGN cluster, we compute the average number of GGN clusters and divide it by the average number expected by chance, computed by randomly generating 200 random ORIs with the same sizes of the real ones. The horizontal axis represents the minimum number of GGN triplets in the clusters, different lines represent different tolerance parameters *L*. Symbols, colors and significances are the same as in Figure 4.

We then analyzed the mean number of GGN clusters of ORIs that contain at least one cluster (Supplementary Figure S9), and compared it with the same number expected by chance. The enrichment, i.e. observed divided by expected clusters per ORI, increases for small *n* and large *L* (Figure 4), which suggests that the selective pressure for finding GGN at ORIs increases in these limits. It is important to note that, despite *C.elegans* has fewer ORIs containing GGN clusters than expected by chance - except for clusters with large *n* and *L* in post-gastrula embryos (Figures 3 and S8) -, these ORIs contain on average significantly more clusters than expected.

Finally, only a small fraction of the GGN clusters overlap with ORIs (Supplementary Figure S10), but their number is much larger than expected by chance except for *D. melanogaster* and *C. elegans* pre-gastrula embryos (Supplementary Figure S11).

In conclusion, ORIs and GGN clusters are significantly associated to each other for different organisms and experimental techniques, in at least one direction and at least for large *n* and *L*, except for *C.elegans* pre-gastrula embryos, but GGN clusters are neither necessary nor sufficient for ORI activation.

### GGN clusters form G-quadruplexes

DNA sequences with tracts of three or more G are prone to fold into G-quadruplex structures (G4) (Bochman et al. 2012). However, this propensity is not so well established for tracts of only two G. We addressed this issue experimentally, by using circular dichroism (CD) and NMR of several oligonucleotides containing repeats of GGA, GGT and GGC triplets. We excluded PolyG sequences since they have a strong tendency to aggregate. The exchangeable proton regions of the 1H NMR spectra of oligonucleotides with eight consecutive repeats of GGA and GGT exhibit sharp imino signals in the range 10.5-12.0 ppm (Fig. 5A). This is consistent with the formation of G4s, as previously reported in the case of some sequences containing GGA repeats (Matsugami et al., 2001). In the case of GGC repeats, additional signals arise around 13 ppm, indicating the formation of G:C Watson-Crick base pairs. These spectra suggest the formation of multiple structures, most probably G4 and B-DNA, in a dynamical equilibrium. In the case of GGA and GGT repeats, the CD spectra with a single maximum around 260 nm indicate the formation of G4 with a parallel topology (Fig. 5B), confirming the NMR data. G4s formed by eight GGN triplets present a high thermal stability, as shown by CD monitored melting experiments (Figure 5C), and NMR spectra recorded at different temperatures (Supplementary Fig. S12). In agreement with previous reports (Renciuk et al., 2009), the CD spectrum of GGC repeats exhibits a negative band around 280 nm, consistent with the presence of B-form DNA. We could not obtain any proper melting curve for this repeat, in agreement with the formation of a mix of multiple structures. Nevertheless, NMR melting experiments indicate that the structures formed by GGC repeats are less stable than those formed by GGA and GGT repeats. These results were obtained for DNA oligonucleotides containing eight triplets, but they can be extrapolated to different numbers of triplets, as shown by NMR experiments recorded for oligonucleotides with four 4 and 12 triplets (see Supplementary Fig. S13).

**Figure 5:**
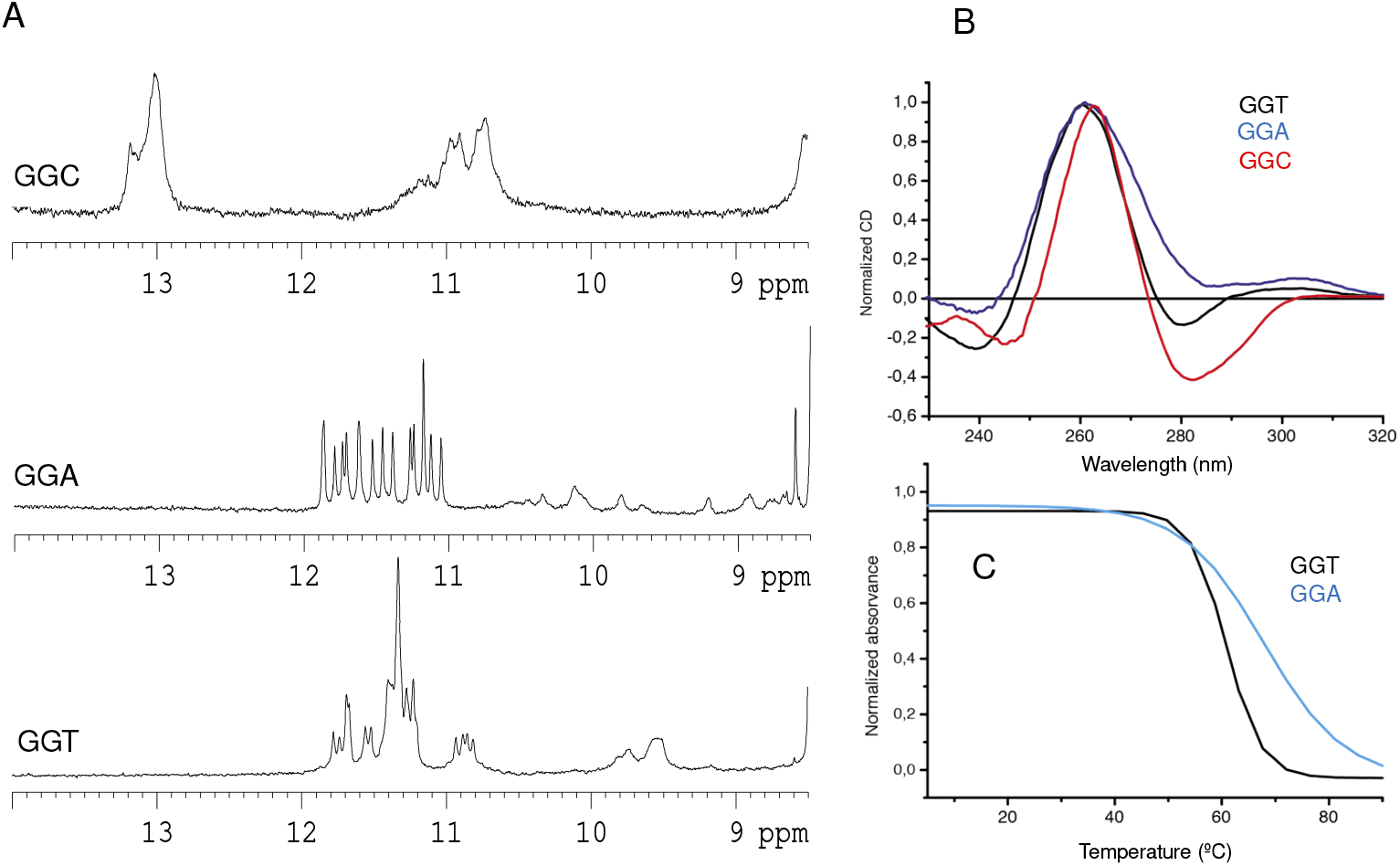
A: Imino proton regions of the NMR spectra of DNA oligonucleotides containing eight GGT, GGA and GGC repeats. B: Circular Dichroism (CD) spectra. C: CD monitored melting curves of eight GGT and GGA, Tm values are 60.9 and 68.5 C, respectively (buffer conditions: 25 mM potassium phosphate, T= 5 C, pH 7).

Interestingly, A is the most frequent base found in the third position of GGN motifs in all studied genomes (except in *L. major*), followed by T (except in *L. major* and *D. melanogaster*). G is the least frequent base at third position of GGN, again with the exception of *L.major* (Supplementary Figure S14). Thus, the observed frequencies recapitulate the ranking of stability of GGN motifs, suggesting that there could be a selective pressure favoring GGN motifs that form stable G4s, although the sequence correlations observed in these regions make it difficult to compare the frequencies with the expectation based on mutational processes that consider both GC-skew and repeated triplets.

### Trinucleotide clusters probably arise through triplet expansion

We then investigated how GGN triplets could arise in evolution. To address this, we measured the propensity to find a GGN triplet at distance *d* from another GGN triplet (Figure 6, black circles, see Methods for details). In all genomes analyzed here, the propensities are positive at small distance, indicating that the presence of a GGN triplet favors the presence of another one at short distance. This occurs for all types of triplets XYN, where X and Y are given and N is any nucleotide, not only GGN (Supplementary Figure S15), suggesting that the mechanism at play is mutational and not selection of any particular triplet. Moreover, in the genomes of *D. melanogaster*, *L. major* and *A. thaliana* GGN triplets show a clear preference for separations equal to an integer number of triplets, as evidenced by the peaks of the propensity at multiples of three bases. Propensities rapidly decay to negative values, suggestive of deletions that take place in highly repetitive regions with many consecutive G nucleotides.

**Figure 6:**
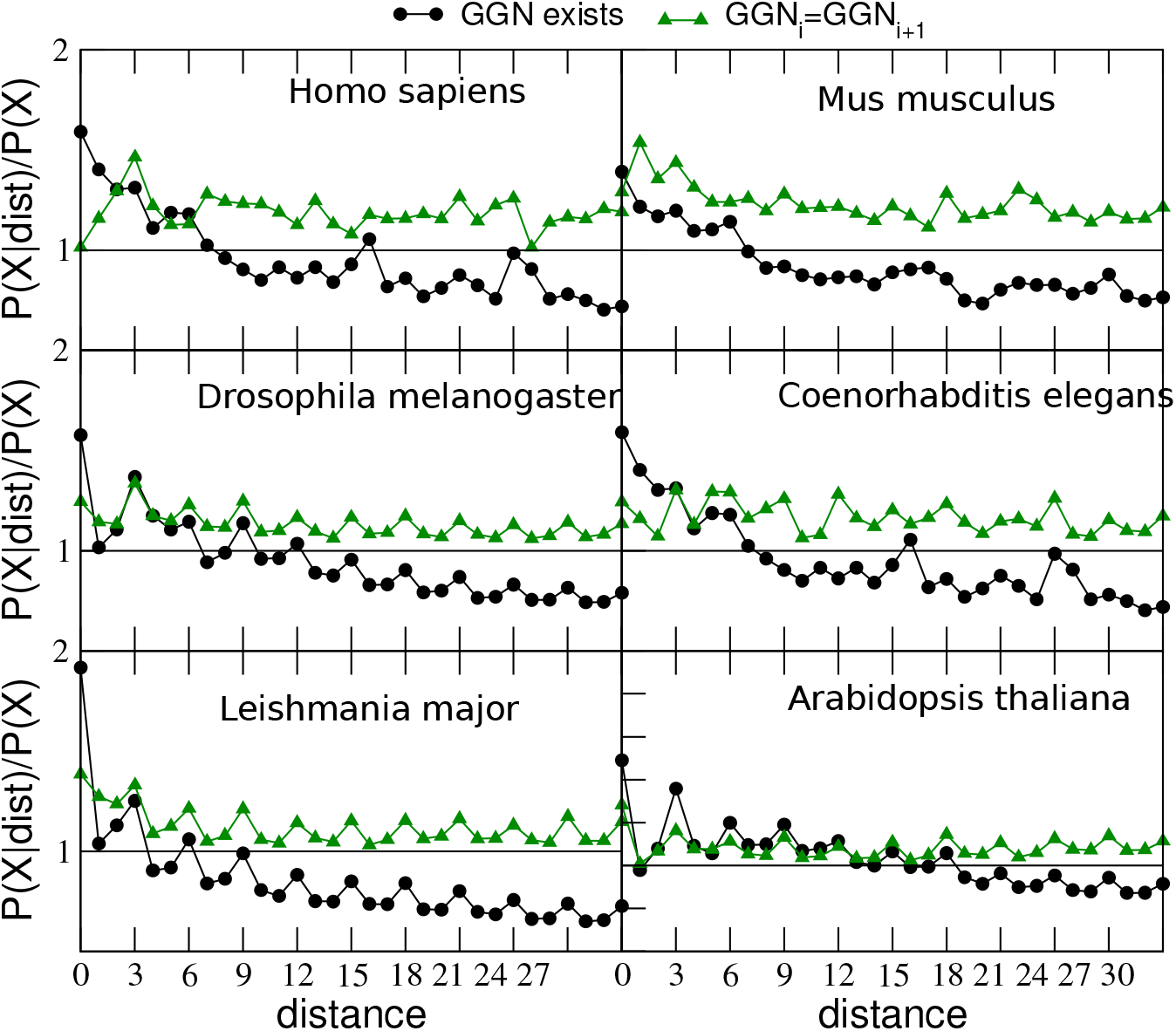
Conditional probability to find a GGN triplet at distance d (in nucleotides) from another GGN triplet (black circles) and that two consecutive GGN triplets at distance d (in nucleotides) have the same third nucleotide (green triangles) divided by the same probability without any condition on the distance.

We also measured the propensity that two consecutive GGN triplets at distance *d* have exactly the same third nucleotide N (Figure 6, green triangles). Again, this propensity is positive in all cases. The genomes of *D. melanogaster*, *L. major* and *A. thaliana* and, less clearly *C. elegans,* show peaks at multiples of three bases, once again consistent with the repetition of an integer number of triplets. Altogether, this data supports the hypothesis that clusters of triplets arise through a mutational mechanism of triplet expansion during genome replication. Again, these properties are not exclusive of GGN triplets but are shared by many types of XYN triplets (Supplementary Figure S16), suggesting that the expansion of short repeats is a pervasive mutational mechanism in eukaryotic genomes.

### GGN clusters associate with precisely positioned labile nucleosomes and with the replication protein CDC6, and locate in the proximity of nucleosome-depleted regions

Fenouil et al. (2012) proposed that G4 structures participate in transcription initiation and ORI specification because they favor the formation of nucleosome-depleted regions (NDR). To test this hypothesis, we examined the relation between GGN clusters and transcript density, nucleosome occupancy and histone variants in the genome of *A. thaliana*. We computed metaplots with respect to the midpoints of the GGN clusters defined with *L ≤ 3* (distance between consecutive GGN) and various values of the minimum number *n* of GGN motifs in the cluster. We oriented the metaplots in the 5’-3’ direction of the longest transcript that overlaps with the GGN, if any.

In contrast to Fenouil et al.’s hypothesis, but consistent with the notion that G+C rich regions have high nucleosome occupancy (Tillo and Hughes, 2009), we found that GGN clusters strongly associate with nucleosomes, as shown by the occupancies of the histone H3 variants H3.3 and H3.1 (Figure 7A and 7B, respectively). The association increases for clusters containing more GGN motifs and it is stronger for nucleosomes that contain the more labile histone variant H3.3. Interestingly, clusters with 8 or more GGN show an adjacent minimum of H3.1 occupancy, i.e. a NDR at ∼1kb from the GGN motif (Figure 7B). These NDR are often associated with the presence of AAN clusters. For instance, we found that clusters of at least 6 consecutive AAN triplets are present at ≤1kb from more than 20% of GGN clusters (Figure 7F), and they strongly favor NDR formation (Fig. 7E). We conclude that these tandem repeats of GGN and AAN triplets, which are common sequence motifs, are associated with characteristic patterns of nucleosome occupancy.

**Figure 7:**
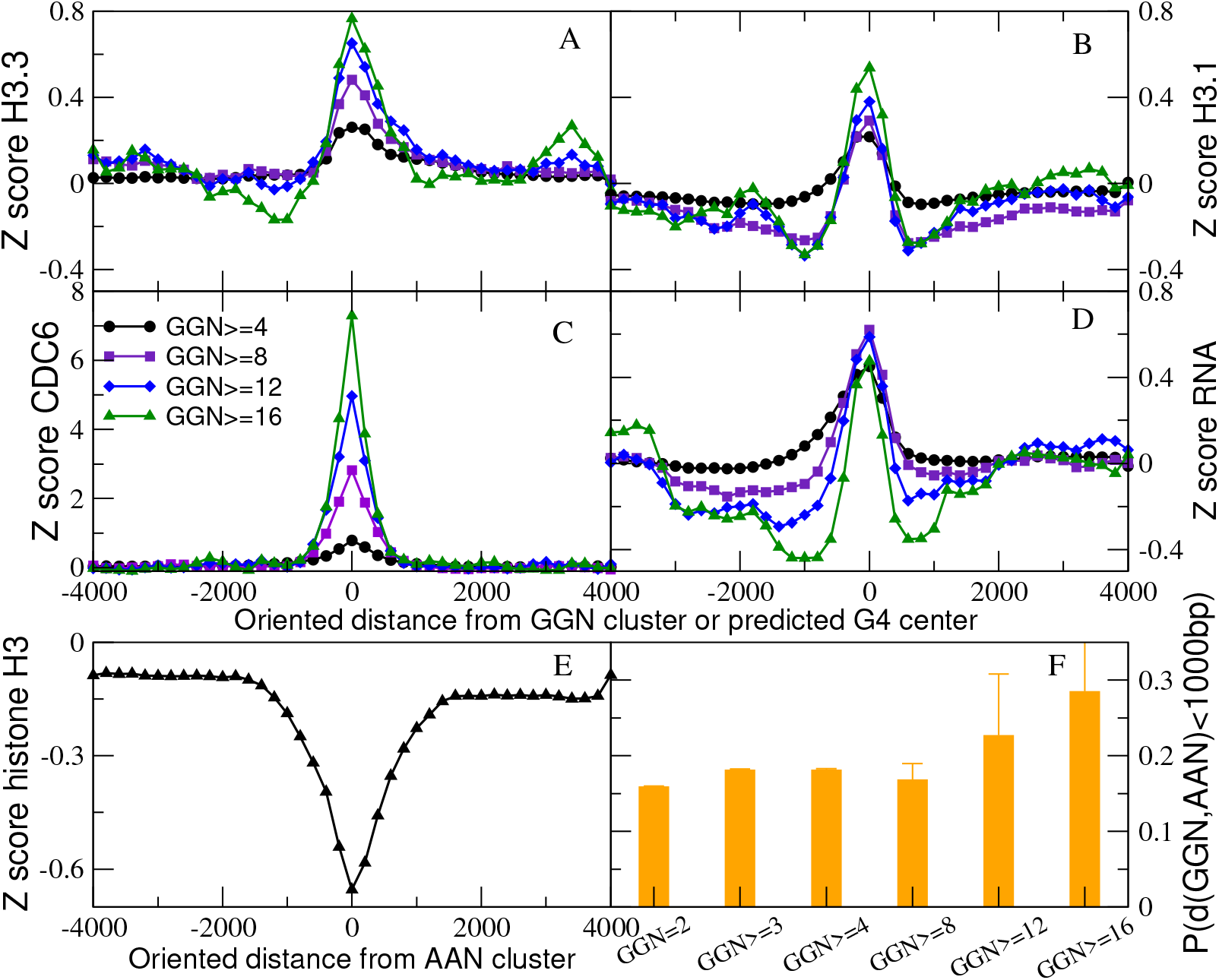
Average Z scores of several features of the *A. thaliana* genome relative to the distance from the center of GGN clusters with *L*=3 and the indicated number of GGN motifs (A-D) and clusters with 6 or more AAN motifs (E). The Metaplots are oriented with the direction of the longer overlapping transcript. Z-scores of the abundances of histone variant H3.3 (A) and H3.1 (B) (GEO accession GSE34840; Stroud et al. 2012); the cell division control protein 6 (CDC6) (C) and RNA abundances (D). (E) Total histone H3 content indicative of nucleosome occupancy (GEO accession GPL10911/GPL10918/GPL11005; Roudier et al. 2011) relative to the center of AAN clusters. (F) Probability to find an AAN cluster at less than 1kb from the center of a GGN cluster, for various lengths of GGN motifs.

We also found that GGN clusters are strongly associated with transcribed regions, since the midpoint of the GGN cluster presents a sharp maximum of transcript density (Fig. 7D). To get more insight on this association, we investigated the distribution of the GGN clusters across the chromatin states determined by Sequeira-Mendes et al. (2014) for the *A. thaliana* genome. Weak clusters with GGN≥4 and *L*≤7 are preferentially associated with the TSS (state 1, Supplementary Figure S17, black bars), but also with GC-rich heterochromatin (state 9). Strong clusters with large n tend to occur in Polycomb-repressed chromatin (state 5), but also with proximal-promoter regions (state 2); for instance, 49% of strong GGN clusters with *n*≥16 and *L*≤3 occur in state 5, and 19% occur in state 2, which is also marked by Polycomb (Supplementary Figure S17, green bars).

Most interestingly, we found a very strong association between the length of the GGN clusters and the pre-replication complex protein CDC6, with Z-score up to 8 (Figure 7C). This association may provide a mechanistic explanation of the association between GGN clusters and ORIs.

### GGN clusters at ORIs and transcriptional start sites

The local landscape of nucleosome occupancy at GGN clusters resembles the nucleosome architecture found at ORIs in several systems (Lombraña et al., 2013; Cayrou et al., 2015; Eaton et al. 2020; Li et al. 2022). In order to address the relationship of nucleosome organization with GGN clusters, TSS, and ORIs, we focus again in the A. thaliana genome, and analyze in detail histone occupancy as well as the location of the replication initiator protein CDC6 at different genomic locations (Figure 8). TSS are marked by an immediately upstream NDR flanked by well-positioned nucleosomes, with a maximum of nucleosome occupancy 500bp downstream (Fig. 8A,B) and enriched in nucleosomes that contain the labile histones H3.3 (Fig. 8B) and H2AZ (Fig. 8C). Importantly, we detected an asymmetry between T and A (AT skew; Fig. 8D) at TSS, likely due to asymmetric mutational processes (Touchon et al., 2004), which may be in part responsible for the NDR.

**Figure 8:**
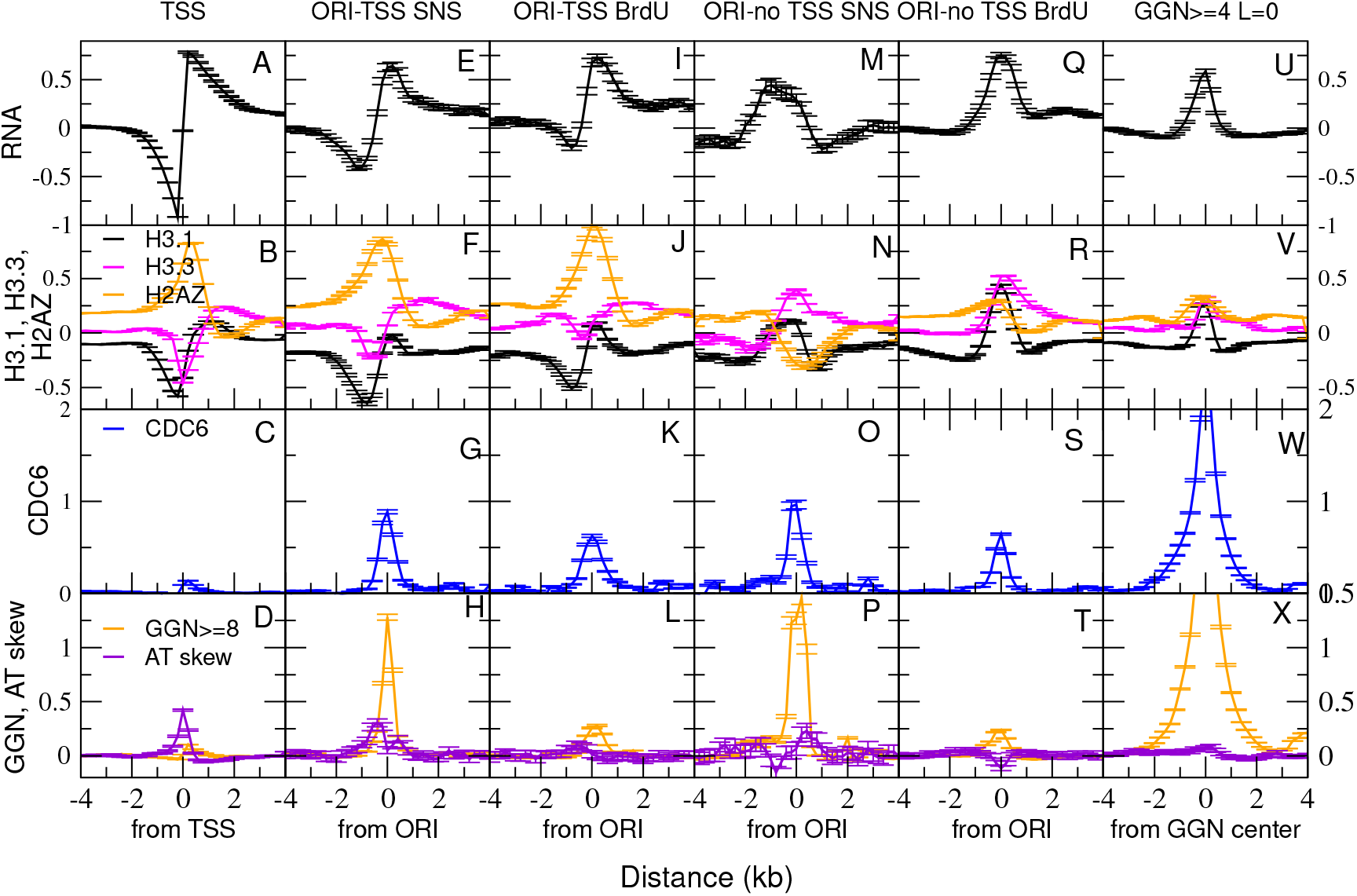
Average values of *A. thaliana* genomic features relative to the distance from specific genomic locations. First column (A-D): distance from transcription starting sites (TSS). Second and third column: distance from the midpoints of ORIs associated with TSS and detected by SNS-seq (second column E-H), or by BrdU-seq (third column, I-L). Fourth and fifth column: distance from ORIs at more than 1kb from TSS and detected by SNS-seq (fourth column M-P) or detected with BrdU-seq (fifth column, Q-T). Sixth column (U-X): distance from the center of GGN clusters with 8 or more GGN separated by 3 or less bases. The y-axis is in unit of Z score with respect to the mean and standard deviation across the entire genome. First row: RNA expression. Second row: relative occupancies of histone variants H3.3, H3.1 and H2AZ. Third row: occupancy of the replication protein CDC6, indicative of ORI activation. Fourth row: AT skew and GGN clusters. All metaplots are oriented in the 5’-3’ direction of the longest transcript found in the plotted region, if any is present. We used the BrdU-ORIs determined by Costas et al. (2011) and the SNS-seq ORIs determined by Sequeira-Mendes et al. (2019).

We next centered the histones signal at ORIs oriented in the direction of overlapping transcripts, and found a remarkably similar organization regardless of the experimental methodology used for ORI mapping (Fig. 8E-H for SNS-ORIs and Fig. 8I-L for BrdU-ORIs). As an important control, the replication initiator protein CDC6 was clearly associated with both sets of ORIs (Fig. 8G,K), but not when all TSS are considered (Fig. 8C). The main difference is that ORIs determined by BrdU labeling present lower GGN Z-scores (Fig. 8H,L), likely because they are more AT-rich. ORIs that are far from the TSS show overall similar features in terms of nucleosome occupancy, histone H3.3 enrichment, association with CDC6 and GGN scores, that are higher for ORIs determined by SNS-seq approaches (Fig. 8M-T). Nevertheless, there are some interesting differences. The labile histone H2AZ is depleted (Fig. 8N) or low (Fig.8R), suggesting that the role of this histone is more relevant for transcription than for DNA replication, and the AT skew is weaker for non-TSS ORIs than for ORIs associated with the TSS (compare Fig. 8H with 8L and 8P with 8T), also suggesting its association with transcription (Fig.8D).

Finally, we centered the metaplots at GGN clusters (Fig.8U-X). As reported above, the association of GGN clusters with the replication protein CDC6 is the highest of all associations found (Z score >2 even for 4 GGN, Fig. 8W) and it increases with the number of GGN in the cluster (Fig. 7C), while the AT skew is not associated with GGN clusters (Fig.8X).

These results lead us to propose the following model of chromatin structure at ORIs (Fig 9):

**Figure 9:**
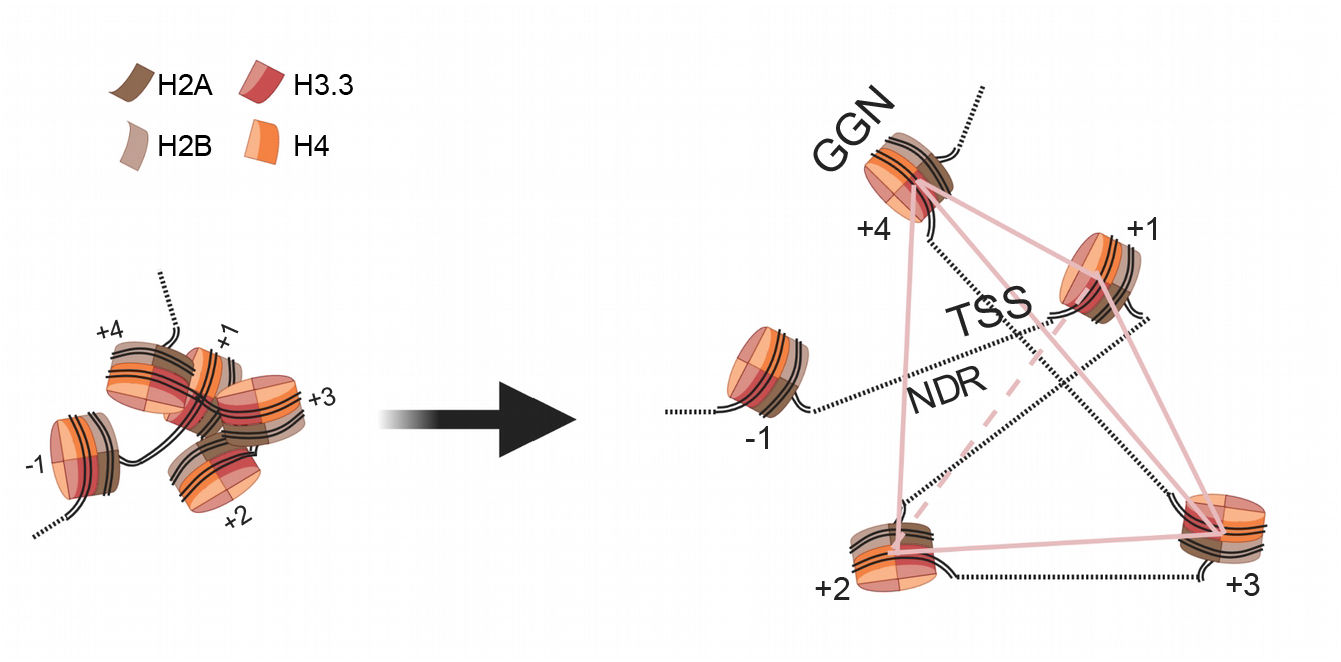
Proposed model of the chromatin structure at ORIs associated with TSS

DNA replication starts at the location of a positioned nucleosome (nucleosome +4), associated with a GGN cluster in *Arabidopsis* and other genomes, and an NDR is present 1kb upstream with the respect to the direction of transcription (for ORIs associated with TSS, Fig. 8F,J) or also downstream (Fig. 8N,R), altogether comprising four nucleosomes. Based on known properties of the chromatin secondary structure (Ohno et al., 2019, see Discussion), we propose a model that assumes that the nucleosome + 4 and the NDR are spatially close.

## Discussion

### GGN triplets: an ubiquitous motif at eukaryotic ORIs

We previously reported that ORIs in the *A. thaliana* genome are enriched in clusters of repeated GGN triplets (Sequeira-Mendes et al., 2019). Here we extended our analysis to a total of six model systems and found that in five out of six genomes, more than 50% ORIs contain a stretch of at least 4 consecutive GGN motifs separated by at most 3 bases, significantly more than expected by chance, reaching 90-95% for the mouse, human and *Leishmania major* ORIs.

Our results are consistent with the notion that ORIs in the genomes of several eukaryotic organisms associate with GC-rich regions (Delgado et al., 1998; Sequeira-Mendes et al., 2009; Costas et al., 2011; Cayrou et al., 2015; Comoglio et al., 2015; Lombraña et al., 2016; Pourkarimi et al., 2016; Rodriguez-Martinez et al., 2017; Vergara et al., 2017; Sequeira-Mendes et al., 2019). They are also consistent with the finding that ORIs tend to be associated with predicted G4 secondary structures of single-strand DNA (Cayrou et al., 2012; Picard et al. 2014; Castillo-Bosch et al., 2014; Prioleau et al., 2017; Akerman et al., 2020). However, it was proposed that these associations may arise in part from experimental bias. Many of the above studies determined ORIs by sequencing short nascent strand after enrichment by λ-exonuclease digestion, which has hindered efficiency at GC-rich sequences. Foulk et al. (2015) argued that this protocol creates a bias towards SNS that are GC-rich. This bias can be reduced by adopting a proper control, also subject to λ-exo digestion, and by carrying out λ-exo treatment under optimal enzyme/substrate conditions (Rodriguez-Martinez et al., 2017; Akerman et al. 2020). Likewise in the *A. thaliana* ORIs that we previously examined (Sequeira-Mendes et al. 2019), which also display a strong correlation between occurrence of GGN motifs and CDC6 binding sites, as expected for genuine replication initiation sites. To address the possible biases due to the experimental methods, here we considered ORI sets obtained through different approaches. In particular, we analyzed in high detail ORIs identified in *A. thaliana* cultured cells through labelling with the thymidine analog BrdU (Costas et al. 2011), a methodology that does not rely on λ-exo treatment and presents the opposite bias towards A+T rich sequences. We did not find any ORI detected with the BrdU method with very strong GGN clusters (20 GGN without any separation, *L*=0, Figure 4), which we interpret as an evidence of the A+T bias of the BrdU method. Nevertheless, the BrdU ORIs identified were also enriched in GGN clusters (Figures 2-4) and show similar properties to the λ-exo-determined ORIs (Figure 8). Therefore, the putative λ-exo bias cannot fully explain the association between ORIs and GC-rich sequences, although the comparison between the two sets of *A. thaliana* ORIs indicates that this bias and the AT bias of the BrdU approach do play a quantitative role. Moreover, the initiation zones of *C. elegans* post-gastrula embryos and human cells determined by Okazaki fragment sequencing (Pourkarimi et al., 2016; Petryk et al. 2016), a methodology that do not involve λ-exo digestion, also show a significant association with GGN clusters. However, OK-seq mapping yields low spatial resolution (the detected ORIs are very broad, Fig.1), which may explain the flat enrichment profiles. Of note, the high-resolution ORIs of *H. sapiens* determined by the Ini-seq method (Langley et al. 2016), which similarly does not involve λ-exo digestion, are more strongly associated with GGN clusters than human ORIs determined through λ-exo (Picard et al. 2014). Besides the experimental approach used, our results suggest that the analysis of the data also play an important role; in particular, it is strongly advisable to use a normalization control that presents the same bias as the experiment as we did for both *A. thaliana* ORI sets.

### Influence of the developmental stage

One unexpected finding from our work is that the proportion of ORIs associated to GGN motifs increase at later developmental stages (Figures 2-4, S5-S9). This occurs in *C. elegans* post-gastrula *vs* pre-gastrula embryos; in *D. melanogaster* Schneider 2 (S2) cells derived from late stage embryos *vs* Bg3-CNS cells that consist of interneurons, motor-neurons and glia arising in the third larval stage (https://flybase.org/reports/FBtc0000068.html); in Mouse Embryonic Fibroblast (MEF) vs. Embryonic Stem (ES) cells; and in *A. thaliana* ORIs from 10 day-old seedlings compared with 4 day-old seedlings. In this later case, we previously found that ORI strength, measured as the number of SNS reads with respect to the control, correlates positively with the number of GGN at the ORI (Sequeira-Mendes et al., 2019). This correlation is stronger than with any other indicator that we measured, including GC content, GC skew, the presence of binding sites of the CDC6 replication protein or the amount of transcriptional activity through the ORI region. Consistent with the observed influence of the developmental stage, the rate of increase of ORI strength *per* GGN increase is stronger for ORIs at 10 days of development (0.056±0.001) than at 4 days (0.046±0.002; Supplementary Figure S18). Altogether, these data suggest that the incidence of GGN clusters is higher in ORIs that are active late in development. It has been recently shown that mouse embryos at later developmental stages activate fewer ORIs (Nakatani et al., 2022). Therefore, it is tempting to speculate that these ORIs activated in late developmental stages correspond to the strongest ones and display the highest association with GGN clusters, thus explaining the increased relationship between GGN clusters and ORIs later in development.

### Structural model of nucleosome organization at ORIs

GGN clusters are strongly associated with nucleosomes, in particular those containing the labile histone H3.3 (Figure 7A and B). The association of the histone variant H3.3 with GC-rich repetitive sequences has also been noted in *Plasmodium falciparum* chromatin (Fraschka et al., 2016). While the association between H3.3 and GGN clusters is general, in *A. thaliana* we observed a specific association between the labile histone H2A.Z and TSS and ORIs associated with TSS (Figure 8C, 8G and 8K) but not for ORIs at other locations (Figure 8N and 8R). Nucleosomes with both labile histone variants mark the position of the origin recognition complex (ORC) in human and mouse cells (Lombraña et al., 2013; Cayrou et al., 2015). Since in *A. thaliana* the variant histone H2A.Z is common at TSS and ORIs that associate with them, our findings suggest that this histone variant is less important for ORI specification in this organism.

The association between GGN sequences and nucleosomes is not surprising, because the G+C content is known to be one of the main determinants of nucleosome occupancy (Tillo and Hughes, 2009). Some years ago, Arneodo and coworkers developed a model of nucleosome positioning (Vaillant et al., 2007; Milani et al., 2009), which computes the energy cost of wrapping a DNA sequence around the nucleosome based on the rotational preferences of trinucleotides (Satchwell et al., 1986; Goodsell and Dickerson 1994). Applying this model to repeated GGN triplets (i.e. trinucleotides of GGN, GNG and NGG), the roll angle per base (Dickerson, 1989) results in 2.34 degrees for GGG; 2.25 for GGT; 2.90 for GGC; and 1.88 for GGA, thus on average 2.39 degrees per base pair for a mix of GGN triplets. This value is very close to the average roll per base pair of the DNA sequence wrapped around the nucleosome (2.47 degrees), which implies that these sequences can wrap around the nucleosome with very little energetic cost, consistent with the high propensity of GGN clusters to localize within nucleosomes. Conversely, for tandem repeats of AAN (consisting of the trinucleotides AAN, ANA and NAA), the average roll per base pair is 0.91 and the corresponding energy cost for nucleosome binding is high, which rationalizes the finding that AAN clusters tend to be associated with NDRs (Figure 7E). Interestingly, we also observed that GGN clusters are often located 0.5-1.5kbp downstream of NDRs (with respect to the transcript direction; Figure 7B), which are often associated with AAN clusters (Figure 7F). Taken together, these results suggest a simple code for nucleosome positioning that can easily evolve through triplet repeats.

TSS and ORIs share a similar organization (Figure 8), namely they are located between an upstream NDR, presumably favored by the large AT content of non-transcribed regions and the AT skew produced by transcription, and a downstream region of high nucleosome occupancy enriched in the labile histone H3.3. The similarity between TSS and ORIs is not surprising, because almost 80% *A. thaliana* ORIs are located at less than 1kb from TSS. We hypothesize that this might occur because both share similar requirements for chromatin accessibility and regulation. The pattern of nucleosome occupancy observed at GGN clusters resembles that of ORIs and TSS, which may partly explain their association.

In *A. thaliana*, the NDR and the stable nucleosome associated with the GGN cluster are separated by approximately 1kb (Figures 7B and 8V). Despite this distance in the linear scale, we propose that the NDR and the well-positioned nucleosome could be in close spatial proximity (Fig 9). This model is inspired by a recent study that proposed two types of chromatin secondary structure (Ohno et al., 2019): (i), alpha, in which four adjacent nucleosomes are disposed at the vertices of a tetrahedron; and (ii), beta, in which they sit at the four vertices of a rhombus (not necessarily planar). 90% of TSS present the beta structure, while gene bodies prefer the alpha structure. The study of Ohno et al. (2019) show two properties that are crucial for our interpretation: i) The TSS is proximal in sequence to an NDR (Fig.2B of Ohno et al. 2019), as we also observe in *A. thaliana* (Fig.7). ii) The nucleosome adjacent to the TSS is in close spatial proximity with the one located four nucleosomes away along the chromatin fiber in the direction of transcription (Fig.S4I of Ohno et al. 2019). We assume that this nucleosome coincides with the stable nucleosome associated with the GGN cluster and the ORI. Building on these findings and on our data, we propose a structural model of nucleosome organization at ORIs associated with TSS in which the spatial proximity between the ORI and the NDR allows physical interactions between the proteins bound to these regions. The model in Figure 9 shows 5 nucleosomes where the TSS and the NDR are located between the −1 and +1 nucleosome of the beta structure and the ORI sits at the +4 nucleosome, which adopts an alpha structure. This proposed organization may be stabilized in part through electrostatic interactions between the naked DNA at the NDR and the stable nucleosome at the GGN cluster. A similar organization may also apply to other regions that favor stable nucleosomes, such as CpG islands, that play the double role of promoters and ORIs in mammalian cells (Delgado et al., 1998; Sequeira-Mendes et al., 2009).

It attracted our attention the parallelism between the above described nucleosome organization and the nucleosome organization of ORIs in the yeast *Saccharomyces cerevisiae* (Eaton et al., 2010). *S. cerevisiae* ORIs contain AT-rich autonomous replicating sequences (ARS), to which the replication initiator protein ORC binds. The 11-base long consensus sequence of the ARS (ACS), consists of two stretches of three or four T separated by three bases, so this sequence tends to avoid nucleosomes (its complementary sequence is analogous to an AAN cluster). Nevertheless, ACS are not sufficient for defining an active ORI. Several thousand ACS were identified in the yeast genome by bioinformatics analysis, but only 250 to 400 of them may act as ORIs. Eaton et al. (2010) showed that ACS at active ORIs are contiguous to other T-rich or A-rich sequences that create an NDR, and they are close to precisely positioned nucleosomes. A recent report strengthened these results, supporting a dual model in which ORC either binds to consensus ACS elements in the *S. cerevisiae* genome or binds to nucleosomes near an NDR (Li et al. 2022). Therefore, the nucleosome organization of *S. cerevisiae* ORIs resembles the one proposed here, emphasizing that ORI organization at the nucleosome level is more conserved than previously thought (Lombraña et al., 2013). An important difference between *S. cerevisiae* and *A. thaliana* ORIs is that well-positioned nucleosomes in yeast are adjacent to the NDR, while in *Arabidopsis* they are far in sequence (Figure 8F,J) but, as we propose, they may be proximal in space (Figure 9).

### Association with the pre-replication protein CDC6

Our analysis suggests that GGN clusters, besides favoring the necessary nucleosome organization, also favor the binding of the pre-replication protein CDC6, which plays an essential role in ORI specification. The larger the number of GGN triplets in the cluster, the stronger its association with the CDC6 protein (Figure 7C). The Z score of this association is the strongest one that we detected in this study.

Furthermore, our analysis suggests that GGN clusters have a dual role in ORI specification, not only favoring the necessary nucleosome organization but also in the binding of the pre-replication protein CDC6 (Figure 7C).

### G4 and their possible role in ORI function

Four consecutive GGN motifs can theoretically form G4 structures with two tetrads, as experimentally demonstrated in some systems (Macaya et al., 1993, Saccà et al., 2005). Indeed, the stability of a G4 structure is largest for tetrads separated by loops of length 1, such as those in consecutive GGN motifs (Chen and Yang 2012). Stabilizing interactions between distinct G4 structures have been observed in the human hTERT promoter (Palumbo et al., 2009), suggesting the possibility of synergies between the large numbers of GGN motifs that we observed in several ORIs (Figure 4). Our CD and NMR spectroscopy studies showed that oligonucleotides composed of at least four repetitions of GGA, GGT and GGC triplets tend to form G4 in vitro, mixed with G:C Watson-Crick base pairs in the case of GGC (Figure 5). In a recent paper, Kejnovská et al. (2021) investigated with CD and NMR spectroscopy the ability of the DNA sequence AATG_2_TG_2_T_3_G_2_TG_2_TAA, containing four GGT triplets, to form an intramolecular parallel-stranded G4. Their results suggest that this sequence forms a transiently populated metastable two-tetrad species, but this is not thermodynamically stable unless it is stabilized through interactions with additional tetrads, *i.e.* forming dimers with four G-tetrads. Dimer formation could explain the relatively large linewidths observed in our NMR spectra of (GGT)4 compared to the DNA constructs with more repeats (see Supplementary Figure S13). In any case, the stability of systems containing eight GGT triplets is consistent with our results (Fig. 5C), and it confirms that GGN clusters with eight or more triplets form stable parallel G4 structures.

The ranking of G4 stability was GGA > GGT > GGC, with *GGG* predicted to be least stable. Interestingly, in all genomes except in *L. major*, which is the richest in GGN triplets, A and then T are the most frequent bases at the third position of GGN motifs (Supplementary Figure S14). Their frequency at the third position is higher than the genomic frequency, suggesting that the observed enrichment may be due to selection, possibly for G4 stability. This hypothesis predicts stronger selective pressure for clusters with fewer GGN motifs or increasing separation between them, two features that negatively affect G4 stability. Indeed, the enrichment of the number of GGN clusters per ORI defined with respect to the number expected by chance is larger for weaker clusters (few GGN or large L, Figure 4), consistent with possible selection for G4 stability at ORIs.

In recent years, there has been debate about the possible role of G4 structures in ORI specification and activity. These structures form when the DNA is single-stranded after replication or transcription, and they may prevent that these processes start again unless specific enzymes resolve the G4. G4 structures hinder the progression of DNA replication forks in mutants of *C. elegans* and human cells lacking the FANCJ helicase (Kruisselbrink, et al., 2008; London et al., 2008). G4 are also a source of genetic instability (Koole et al. 2014), which may occur when the replication stalls ahead of structured G4, unless the primase-polymerase PrimPol acts to reprime replication (Schiavone et al. 2016). On the other hand, two G4-containing ORIs can drive DNA replication initiation in the chicken DT40 cell line (Valton et al., 2014, Prorok et al., 2019). More recently, Poulet-Benedetti et al. (2023) found that two G4-forming motifs on the same strand are sufficient for replication initiation. According to this study, however, tandem G4s associated with ORIs tend to be located in NDR next to well-positioned nucleosomes, which is different from the GGN clusters studied here that are strongly associated with nucleosomes.

Three main hypothesis have been proposed to explain the role of G4s in ORI specification: (1) G4s may exclude nucleosomes, thereby potentially favoring the formation of pre-replication complexes (Fenouil et al. 2012). (2) G4s may facilitate DNA melting (Valton & Prioleau 2016). (3) G4s may facilitate the recruitment of specific factors involved in the formation of a functional origin (Hoshina et al. 2013; Keller et al. 2014). Hoshina S, Yura K, Teranishi H, Kiyasu N, Tominaga A, Kadoma H, Nakatsuka A, Kunichika T, Obuse C, Waga S. Human origin recognition complex binds preferentially to G-quadruplex-preferable RNA and single-stranded DNA. J Biol Chem. 2013 288:30161-30171. doi: 10.1074/jbc.M113.492504 Keller H, Kiosze K, Sachsenweger J, Haumann S, Ohlenschläger O, Nuutinen T, Syväoja JE, Görlach M, Grosse F, Pospiech H. The intrinsically disordered amino-terminal region of human RecQL4: multiple DNA-binding domains confer annealing, strand exchange and G4 DNA binding. Nucleic Acids Res. 2014 42:12614-27. doi: 10.1093/nar/gku99 Our findings are in contrast with hypothesis 1, and suggest an additional hypothesis: GGN clusters in the double stranded state can localize nucleosomes precisely, with strong preference for the labile histone H3.3. This nucleosome organization is thought to be important for ORI activation (Lombraña et al. 2013), and it may contribute to explain why GGN clusters are so frequently associated with ORIs. The same GGN clusters are able to form G4 structures when they are single stranded, which creates an indirect association between ORIs and G4. The hypothesis (3), i.e. that G4 may favor the recruitment of initiator proteins is consistent with our observation that, at least in *A. thaliana*, strong GGN clusters favor the binding of the pre-replication protein CDC6 (Figure 7C) even more strongly than ORIs. However, our data cannot discern whether this association is due to a physical interaction and whether G4 structures mediate it. The latter hypothesis would be consistent with the recent finding that human Origin Recognition Complex subunit 1 (hORC1), which is homolog of CDC6, preferentially binds to G4 DNA through a disordered stretch of 99 residues (413-511) in which basic and polar residues (arginines, lysines, serines and threonines) perform key interactions with the G4 (Eladi et al. 2023). Thus, in our view, GGN clusters are associated with ORIs mainly because they create a favorable nucleosome organization and, at the same time, they favor CDC6 binding. Furthermore, these structures form when the DNA is single-stranded after replication or transcription, and they may prevent that these processes start again unless specific enzymes resolve the G4. In the case of transcription, this hypothetical effect exists only for G4 that form in the sense strand, which slightly exceed 50% in *Arabidopsis,* although this bias may be due to mutational bias.

### GGN clusters may evolve through cluster expansion

Finally, we investigated the possible evolutionary mechanism that creates GGN clusters. We propose that the enrichment in GGN stems from two genomic properties of ORIs, the high GC content and the GC-skew, which concur to produce one G-rich strand. This GC skew associated with DNA replication was attributed to the mutational asymmetry between the leading and lagging strands (Lobry 1996). However, we found that in *A. thaliana* ORIs the asymmetry between G and C starts approximately 300 bp before the ORI midpoint, which we located as the maximum of the nascent strand score (Sequeira-Mendes et al., 2019). Thus, DNA replication asymmetry cannot be the sole cause of the GC skew of ORIs in this organism. GGN clusters are tandems of quasi-repeats, and a likely mutational mechanism through which they might arise is triplet insertions produced by the slippage of the DNA polymerases, which happens frequently at homo-polymeric tracts. This hypothetical mechanism is supported by the positive propensity to find a GGN triplet at a certain distance from a similar triplet, which presents clear maxima at multiples of 3 bases in the genomes of *L. major*, *D. melanogaster* and *A. thaliana* (Fig. 6). Moreover, the propensity that the third nucleotides N are identical at consecutive GGN triplets also presents maxima at distances of multiples of 3 bases in the same genomes. In our view, GGN clusters at ORIs are then maintained by natural selection that favors correct nucleosome organization and possibly G4 structures.

## Conclusions

Clusters formed by several repeats of GGN triplets are ubiquitous in eukaryotic DNA replication origins (ORIs). Although they are neither necessary nor sufficient for ORI specification, they correlate with ORI activation strength, at least in *A. thaliana*. Furthermore, ORIs containing GGN clusters are more frequently used at later developmental stages, when fewer ORIs fire. We hypothesize that these preferred ORIs are those that are associated with GGN clusters, which appear in many analysis as the most efficient ones.

We presented here NMR and CD experiments that show that single-stranded GGN clusters can form G4 structures *in vitro*. Furthermore, based on a biophysical model of nucleosome positioning (Vaillant et al. 2007), GGN clusters are predicted to have strong propensity to co-localize with nucleosomes, as confirmed by genomic data. The GGN clusters show a high preference for nucleosomes containing the labile histones H3.3, and they have a very strong tendency to co-localize with the pre-replication protein CDC6. Nucleosome depleted regions (NDR), frequently associated with T-rich tracts, occur frequently at 600-1200 bp distance from GGN clusters and ORIs. Based on the properties of chromatin secondary structure (Ohno et al. 2019), we propose a structural model in which NDR are close in space to a well-positioned nucleosome, allowing functional interactions. This model recapitulates the relative positions along the genome of ORIs, TSS and NDR. The proposed local organization of nucleosomes unifies GGN clusters, TSS, yeast ORIs that present a consensus sequence and ORIs of higher eukaryotes.

We suggest that GGN clusters at ORIs arose in evolution through the synergy between robust mutational mechanisms (mutational skews amplified through triplet expansion) and natural selection for a favorable nucleosome organization and for specific binding of replication proteins. In our view, this synergy constitutes an important ingredient of the evolvability of living systems.

## Methods

### Data sets

We downloaded the genomic coordinates of several sets of experimentally determined replication origins of six model genomes, as reported in the Results section and summarized in Supplementary Table 1.

For *Arabidopsis thaliana,* we reanalyzed the data of Ref. (Costas et al. 2011), which were obtained by labeling with BrdU newly synthetized DNA. We reanalyzed this data with our program ZPeaks, the same as we used for calling A. thaliana ORIs from SNS-seq datasets from seedlings (Sequeira-Mendes et al. 2019). In order to take into account that only thymine bases can be labeled with BrdU, we used as a control the number of T bases in each control read, so that T-rich sequences receive high counts both in the experiment and in the control. The resulting set contained 5824 ORIs. It is available in Supplementary Material.

We downloaded the genome sequence of these genomes, for the same release used for the ORI coordinates, respectively: *H. sapiens*: hg19; *M. musculus*: mm10; *D. melanogaster*: dm3; *C. elegans*: ce10; *L. major*: Lmjf6; *A. thaliana*: TAIR10.

### Bioinformatics analysis

We used our own program GGN_cluster in order to retrieve from an input genome the complete list of clusters containing at least *n* GGN triplets, where N is any nucleotide, separated by at most *L* other nucleotides. In general, the program analyzes all possible types of XYN triplets, where XY is a specified di-nucleotide, and prints them in bed files if requested by the user. Moreover, the program may also analyze and print clusters of XY doublets with the same parameters. A XY cluster with *L*=1 consists of XYN clusters with *L*=0 plus consecutive instances of the XY doublet.

The program GGN_cluster also examines statistical properties of XYN and XY clusters for all possible values of X and Y that yield different clusters and given *l*. These properties are: The number of clusters with exactly *n* motifs. ii) The propensity to find a triplet XYN or a doublet XY at distance *d* from another triplet of the same type, i.e. the ratio between the conditional probability to find a motif at distance *d* from an identical motif divided by the unconditioned probability. iii) The probability that two XYN motifs at distance *d* have the same third nucleotide N, as a function of *d*. The resulting curves provide hints on whether consecutive XYN or XY motifs are produced through a process of triplet expansion or doublet expansion.

Finally, we computed the overlap between the lists of GGN motifs and the list of ORIs, both in bed format, through our in-house program Compare_peaks that also computes the probability to observe the same overlap by chance. To do so, the program randomly draws 200 samples of the same number of genomic sequences with exactly the same size and random coordinates, both for the first and the second compared list, computes the overlap with the other list and computes the probability that the observed overlap is larger than the random one.

### NMR experiments

DNA samples for NMR experiments were suspended in 500 µL in H2O/D2O 9: 1 in 25mM potassium phosphate, pH 7. NMR spectra were acquired in Bruker Avance spectrometers operating at 600, MHz, and processed with Topspin software. Water suppression was achieved by including a WATERGATE module in the pulse sequence prior to acquisition.

## Availability of data and materials

The software programs and the datasets supporting the conclusions of this article are available under GNU licence in the github repositories https://github.com/ugobas/GGN_clusters and https://github.com/ugobas/Compare_peaks

## Acknowledgements

This work was supported by the Spanish government and the Agencia Estatal de Investigación (AEI, 10.13039.501100011033) under grants BIO2016-79043-P (MEC/FEDER) and PID2019-109041GB-C22 (AEI) to U. Bastolla, PID2020-116620GB-I00 to C. Gonzalez, PID2019-105949GB-I00 (AEI) to M. Gómez and PID2021-123319NB-I00 (MICIN/FEDER) to C. Gutierrez, and by the EU under grant ERC-2018-AdG-833617 to C. Gutierrez. Research at the CBMSO is supported by institutional grants from Banco Santander and Fundación Ramón Areces. We acknowledge the ‘‘Manuel Rico’’ NMR laboratory (LMR), a node of the ICTS R-LRB for the use of the NMR instruments, and we gratefully thank Ramón Peiró for technical help.

